# Separation, isolation, and membrane lipid profiling of selectively vulnerable and resilient neurons

**DOI:** 10.64898/2026.10.06.757005

**Authors:** Michael Sadek, Midori Yokomizo, Emily Williams, Natalia Wieckiewicz, Desmond Owusu Kwarteng, Sudeshna Das, Rachel E Bennett, Teresa Gomez-Isla, Masato Maesako

**Affiliations:** Harvard Medical School, 25 Shattuck Street, Boston, MA 02115, USA; Department of Neurology, Massachusetts General Hospital, 114, 16th street, Charlestown, MA 02129, USA

**Author notes:** Correspondence: Masato Maesako, Ph.D. These authors equally contributed (Michael Sadek, Midori Yokomizo).

**Keywords:** Neuronal vulnerability and resilience, membrane integrity, time-lapse imaging, FACS, lipidomic analysis

## Abstract

Certain neurons are more susceptible to pathogenic stressors and degenerate, while others remain resilient in neurodegenerative diseases. However, the mechanisms underlying this cell-to-cell heterogeneity in neuronal vulnerability remain unclear, primarily due to challenges in isolating vulnerable and resilient neurons to characterize their properties. In this study, we report the development of a novel cell-labeling method utilizing two cell-impermeable dyes to identify vulnerable and resilient neurons in culture, which are subsequently separated and isolated via fluorescence-activated cell sorting (FACS). As a proof-of-concept of a mechanistic study, we conducted lipidomic analysis to determine whether the lipid composition of membranes differs between vulnerable and resilient neurons to oxidative stress. This simple but unique method sets the groundwork for investigating the molecular underpinnings of selective neuronal vulnerability, which could inform new diagnostic strategies and therapeutic targets for neurodegenerative diseases.

## Introduction

Neuronal loss is the critical pathological event in neurodegenerative diseases such as Alzheimer’s Disease (AD). This is driven by multiple internal and external stressors, including the accumulation of misfolded proteins, neuroinflammation, excitotoxicity, and oxidative stress, and it most likely occurs through both programmed signaling pathways and non-programmed cell death (Andreone et al., 2020; Moujalled et al., 2021). One fundamental aspect of cell death in neurodegenerative diseases is selective neuronal vulnerability, in which distinct cell populations exhibit different susceptibilities to pathogenic processes and degeneration (Fu et al., 2018; Kampmann 2024). In other words, some neurons undergo cell death in response to pathogenic stressors (i.e., vulnerable neurons), whereas others remain unaffected (i.e., resistant) or sustain damage, recover, and ultimately survive (i.e., resilient). Uncovering the mechanisms and consequences of this selective vulnerability is therefore crucial to our understanding of etiology and pathophysiology, as well as to the identification of potential therapeutic targets and diagnostic markers for neurodegenerative diseases. However, such endeavors have largely been limited by the lack of methodologies to identify vulnerable and resistant/resilient neurons and physically separate them for mechanistic studies.

Single-cell RNA sequencing (scRNA-seq) technology, which enables robust transcriptional profiling of neurons from diseased human tissues and *in vivo* animal models, has been actively used to study neuronal vulnerability. Recent results show that manifestations of selective vulnerability across brain regions may be explained by differences in cell-type specific transcriptional programs that affect various processes such as protein folding, autophagy, apoptosis, mRNA regulation (Saura et al., 2023). To study neuronal vulnerability from a different angle, we previously employed an *in cellulo* model and developed a time-lapse live-cell imaging assay based on the quantitative assessment of membrane integrity (Yokomizo et al., 2025). This unique assay enables us to quantitatively assess, with single-cell resolution, which neurons die in response to toxic insults (e.g., oxidative stress, excitotoxicity) and the time at which they die. Using this approach, we reproducibly distinguished vulnerable from resistant and resilient neurons in culture, which has opened a new avenue to study the upstream regulators and downstream consequences of selective neuronal vulnerability. While our assay cannot distinguish whether neurons are unaffected (i.e., resistant) or become damaged, then recover and ultimately survive (i.e., resilient), we refer to this subset of neurons hereafter as simply ‘resilient’ neurons.

In this study, we expand on the assay above and report the development of a novel cell labeling and isolation method, enabling us to differentially color-code vulnerable, resilient, and already dead neurons, and physically separate and isolate them via fluorescence-activated cell sorting (FACS). Since lipids are tightly associated with membrane integrity (Harayama and Riezman, 2018), we performed lipidomic analysis of membrane fractions sorted from vulnerable and resilient neurons to oxidative stress to explore potential differences in their membrane lipid compositions. This study establishes a rigorous experimental framework that enables us to physically separate and isolate functionally validated vulnerable and resilient neurons in culture. Moreover, it demonstrates a proof-of-concept for utilizing this assay to explore factors associated with selective neuronal vulnerability, leading to new therapeutic strategies and improved diagnostic approaches for neurodegenerative diseases.

## Material and Methods

### Reagent

SYTOX™ Green Nucleic Acid Stain (SG) and Propidium iodide (PI) were purchased from Thermo Fisher Scientific (Waltham, MA). 4,4’-dithiodipyridine (DTDP), 4-hydroxynonenal (HNE), Dimethyl sulfoxide (DMSO), Triton X-100, Chloroform, and Methanol were from Sigma-Aldrich (St. Louis, MO). Conduritol B epoxide (CBE) and Miglustat were from MedChemExpress LLC (Monmouth Junction, NJ)

### Neuronal culture

The Papain Dissociation System (Worthington Biochemical Corp, Lakewood, NJ) was used to dissociate primary cortical neurons from CD1 mouse embryos at E14–16 (Charles River Laboratories, Wilmington, MA). Neurons were cultured in Neurobasal medium supplemented with 2% B27, 1% GlutaMAX, and 1% penicillin–streptomycin (Thermo Fisher Scientific) for 6–9 days in vitro (DIV) before experiments. The animal protocol for neuron preparation was approved by the Massachusetts General Hospital Animal Care and Use Committee (2003N000243).

### Sequential cell labeling to identify dead, vulnerable, and resilient neurons

For confocal time-lapse imaging, neurons were incubated with SG at a final concentration of 0.5 μM for 15 min at 37°C under light-restricted conditions. Cultures were then washed to remove extracellular dye. Immediately after washout, PI was added to a final concentration of 5 μg/mL together with 2.5 mM DTDP. DMSO at the corresponding dilution was used as a vehicle control. Right after adding PI and DTDP or vehicle control set as 0 min time point (T = 0 min).

For flow cytometry/FACS, neurons were incubated with SG at a final concentration of 25 nM for 15 min at 37°C under minimal light exposure, then washed with PBS−/−. PI was then added at a final concentration of 1 μg/mL, together with 0.5-1.0 mM DTDP or vehicle, and the mixture was subjected to flow cytometry and/or sorting as described below.

### Time-lapse live-cell imaging

An Olympus FV3000RS Confocal Laser Scanning Microscope (Tokyo, Japan) was used for time-lapse fluorescence imaging. Suitable CO_2_ concentration and heating for live-cell imaging were maintained using a CO_2_/heating unit (Tokai-Hit, Fujinomiya, Japan). The TruFocus Z drift compensation module was used to maintain focus during time-lapse imaging. Images were acquired with an interval of 15 min for a total duration of 12 h. A 10x/0.40NA objective was used for image acquisition. A 488 nm laser was used to excite SG, and emission was detected between 500 and 540 nm. A 561 nm laser was used to excite PI, and its emission was detected between 600 and 680 nm. Laser power and detection parameters were kept constant within each experiment. For some of the experiments, the BioTek Cytation5 cell imaging multimode reader (Agilent Technologies, Santa Clara, CA) was used.

Imaging data were analyzed using ImageJ/Fiji (NIH). Time-lapse stack images were opened, and an image after DTDP + PI addition, i.e., T = 0 min, in the bright-field channel was used to create cytoplasmic regions of interest (ROIs). ROIs were manually drawn on individual neurons in an unbiased manner, and the same ROIs were applied to the corresponding PI channel using the ROI Manager. Mean PI fluorescence intensity (arbitrary units (a.u.)) within each ROI was quantified across the time series using the Time Series Analyzer V2 plugin. Neurons were classified as PI-positive when the ROI mean PI intensity exceeded 250 a.u, and the first time point at which the ROI mean PI intensity crossed this threshold was defined as the time until becoming PI-positive.

### Fluorescence-activated cell sorting (FACS)

Following the sequential labeling described above, neurons were scraped and filtered through a 40 µm cell strainer to remove cell aggregates. Flow cytometry and cell sorting were performed at the MGH Pathology, Flow, and Mass Cytometry Core on the BD FACSMelody (BD Biosciences, Franklin Lakes, NJ) equipped with 3 laser lines and a 100 µm nozzle. Neurons were excited with a 488 nm laser; the emitted fluorescence from SG and PI was detected simultaneously in the 505-559 nm and 567-597 nm emission channels, respectively. 1% Triton X-100 for 10 min incubation was used to define the dead cells gate and confirm quadrant placement. In the gating, debris was excluded by FSC/SSC, singlets were gated by FSC-A vs FSC-H, and populations were classified on a SG vs PI bivariate plot. Baseline-dead neurons were identified as SG-positive at T = 0 (and typically PI-positive after PI addition), vulnerable neurons were identified as SG-negative/PI-positive, and resilient neurons were identified as SG-negative/PI-positive. Controls included unstained cells, SG-only, PI-only, vehicle-treated, DTDP-treated, and fully permeabilized neurons using 1% Triton X-100. The number of sorted neurons and their viability (%) were assessed using the LUNA-FL™ Dual Fluorescence Cell Counter (Logos Biosystems, Annandale, VA) with the Acridine Orange/Propidium Iodide Stain (Logos Biosystems).

### Extraction of membrane fractionation

Sorted neurons were immediately pelleted at 14000 × g for 1 min at 4°C. Cytosolic and membrane fractions were purified using the Subcellular Protein Fractionation Kit for Cultured Cells (Thermo Fisher Scientific) according to the manufacturer’s protocol. For Western blotting, protein concentration was measured using the BCA Protein Assay kit (Thermo Fisher Scientific). After mixing the samples with NuPAGE^TM^ LDS Sample Buffer and Sample Reducing Agent (Thermo Fisher Scientific), the samples were boiled, subjected to SDS-PAGE on NuPAGE^TM^ 4-12% Bis-Tris Protein gels (Thermo Fisher Scientific), and transferred to nitrocellulose membranes (Thermo Fisher Scientific) using the BioRad Wet electroblotting system (BioRad, Hercules, CA). The membranes were incubated with primary antibodies and their corresponding fluorophore-conjugated secondary antibodies, then developed on the LI-COR Odyssey CLx scanner (LI-COR Biosciences, Lincoln, NE). The successful purification was verified using anti-Na+/K+-ATPase (membrane fraction) (RRID:AB_309699, MilliporeSigma, Burlington, MA) and anti-β-tubulin (cytosolic fraction) (RRID:AB_2210545, Cell Signaling Technology, Danvers, MA) antibodies.

### Lipid extraction and LC–MS/MS lipidomic analysis

Membrane fractions from vulnerable and resilient neurons were subjected to methanol: chloroform 2:1 (v/v). After chloroform and water were added, the mixture was agitated and vortexed. The mixture was then centrifuged at 3500 rpm to separate the two phases, and the lower chloroform phase, containing lipids, was collected, evaporated, and stored at -20°C. Lipidomic analysis was performed at the Beth Israel Deaconess Medical Center-Harvard Mass Spectrometry Facility, as described previously (Breitkopf et al., 2017; Lundin et aal., 2025). Lipid fractions were analyzed by high-performance liquid chromatography coupled to tandem mass spectrometry (LC–MS/MS) using the Q Exactive [Plus/HF] Orbitrap HR-LC– MS/MS spectrometer (Thermo Fisher Scientific). Lipid species were identified based on accurate mass, retention time, and fragmentation patterns using LipidSearch software (Thermo Fisher Scientific).

To perform statistical analysis of data from independent experiments, the signal peak area for each lipid group was first measured and normalized by the number of cells within each sample. Lipid abundance was then converted to relative percent composition by dividing each lipid value by the sum of all lipids measured in that experiment. The heatmap was generated using GraphPad. For lipid category analysis, all lipid classes within each category were summed and converted to a percentage of that category. Univariate comparisons between the two groups were performed using multiple paired Wilcoxon tests, with multiple-comparison correction using the Holm-Sidak method. For lipid species analysis, a volcano plot was generated to show species that significantly differed between vulnerable and resilient neurons. Given this was an exploratory analysis, univariate comparisons between the two groups were performed using multiple paired Wilcoxon tests without correction for multiple comparisons. In structural categorization analyses, lipids were grouped by total acyl chain length into Short (<6 carbons), Medium (6-12 carbons), Long (13–21 carbons), and Very Long (>21 carbons) categories. Moreover, saturation profiles were categorized based on the total number of double bonds into Saturated Fatty Acids (SFA; 0 double bonds), Monounsaturated Fatty Acids (MUFA; 1 double bond), and Polyunsaturated Fatty Acids (PUFA; 2 or more double bonds). Lipid classes and species with p < 0.05 were considered significantly altered.

### Statistical analysis

GraphPad Prism 10 (GraphPad Software, La Jolla, CA) was used for statistical analysis. The D’Agostino and Pearson omnibus normality test was used to examine the Gaussian distribution of the data. Unpaired t-test, Mann-Whitney U test, Paired Wilcoxon test, one-way ANOVA, Kruskal-Wallis nonparametric test, and Logrank (Mantel-Cox) test were used to compare the data. At least three independent experiments were performed to ensure the reproducibility of the results. The sample size calculation was based on previous results from our laboratory (imaging: Yokomizo et al., 2025, lipidomics: Lundin et al., 2025) and power calculations.

## Results

### A novel sequential cell-labeling workflow enables the identification of baseline-dead, vulnerable, and resilient neurons to DTDP-induced oxidative stress and degeneration

It is highly presumable that individual neurons exhibit distinct levels of susceptibility to toxicity and cell death. To validate this paradigm experimentally, we previously developed a time-lapse imaging assay in which neurons are challenged with a toxic insult, e.g., oxidative stress, excitotoxicity, in the presence of propidium iodide (PI): a membrane-impermeable fluorescent dye. By longitudinally recording membrane integrity, this assay enabled us to quantitatively assess which neurons die and when death occurs at the single-cell level. We reproducibly detected neurons with distinct levels of vulnerability and uncovered that *γ*-secretase and the endo-lysosome system are upstream regulators (Yokomizo et al., 2025). Of note, we verified that our culture system contains MAP2-positive neurons at a high rate (∼94%) (Lundin et al., 2025). In the present study, we optimized the initial approach in Yokomizo et al., 2025, and established a novel, sequential cell-labeling workflow using two cell-impermeable fluorescence dyes, SYTOX™ Green (SG) and PI. As outlined in **Fig. 1**, neurons were first labeled with SG to identify baseline-dead cells, then washed to remove extracellular dye from the conditioned medium. Neurons were then challenged with 4,4’-dithiodipyridine (DTDP) oxidant in the presence of PI to label newly dying neurons in response to DTDP-induced oxidative stress. In this framework, we hypothesized that baseline-dead neurons are SG-positive prior to stress induction and become double-positive after PI and DTDP incubation. On the other hand, a subset of initially SG-negative live neurons acquires a PI signal over time in the presence of toxic insult, which is classified as ‘vulnerable’ neurons. Neurons that remain SG/PI-negative throughout the observation window are ‘resilient’ neurons **(Fig. 1**: Step 1). We further hypothesized that vulnerable and resilient neurons can be physically separated and isolated using FACS **(Fig. 1**: Step 2). To test these hypotheses, we run the following experiments.

**Figure 1.**
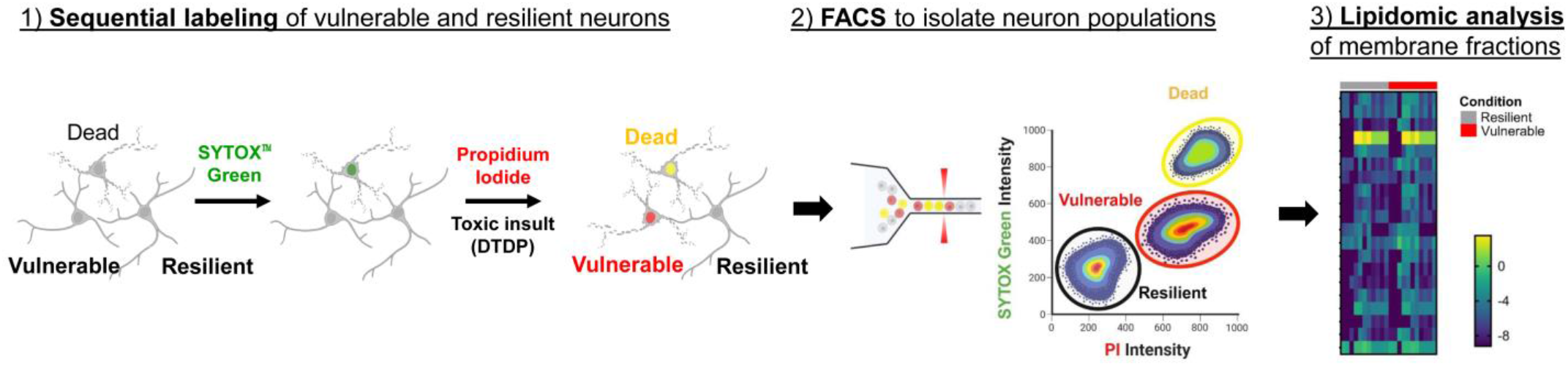
Schematic workflow of identifying and isolating vulnerable vs. resilient neurons to DTDP-induced oxidative stress and degeneration, followed by membrane lipidomics: 1) Cultured neurons are first labeled with SYTOX™ Green (SG) to mark dead neurons (green), followed by addition of PI together with DTDP, an oxidative stressor, such that baseline-dead neurons become SG and PI double-positive (yellow). On the other hand, newly dying neurons (vulnerable neurons) become PI-positive (red), while resilient neurons remain non-fluorescent. 2) Neuronal subpopulations are separated by FACS based on SG and PI fluorescence. 3) Membrane fraction was extracted from sorted vulnerable and resilient populations and subjected to lipidomic analysis.

### Time-lapse imaging validates the dual-dye labeling kinetics and successful distinction of vulnerable and resilient neurons

Time-lapse live imaging was first performed to test our hypotheses. In the experiment, neurons were incubated with 0.5 μM SG for 15 minutes. After washing out the dye from the conditioned medium, the same neurons were co-incubated with 2.5 mM DTDP and 5 μg/mL PI. Time-lapse imaging was conducted to detect SG and PI fluorescence for 2+ hours at every 15-minute intervals from 15 minutes before SG incubation to 120 minutes after DTDP + PI incubation **(Fig. 2A)**. Representative images from time points between -15 and 30 minutes demonstrate that there are neurons that are SG-positive prior to oxidative challenge. As expected, these baseline-dead neurons become double-positive with SG and PI immediately after PI and DTDP addition (yellow). In contrast, some neurons transition from SG/PI double-negative to PI-positive (red) following PI + DTDP exposure, suggesting the existence of ‘vulnerable’ neurons. However, we found that other neurons remain non-fluorescent for a certain period of time, which can be ‘resilient’ neurons **(Fig. 2B)**.

**Figure 2.**
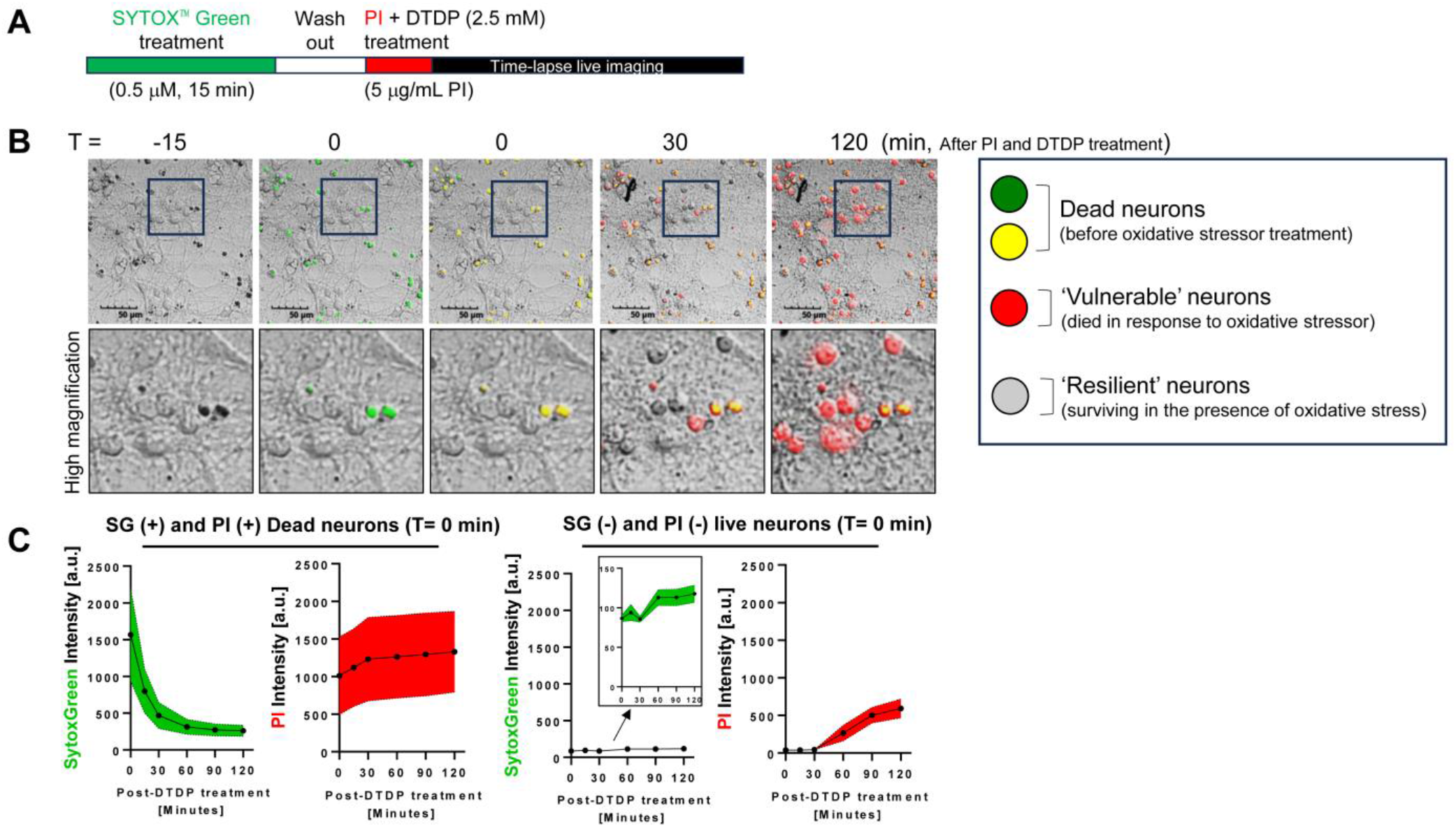
Time-lapse imaging validates the kinetics of sequential SYTOX™ Green and PI labeling: **(A)** Experimental timeline: neurons were treated with 0.5 μM SG for 15 min, washed to remove extracellular dye, and then treated with 5 μg/mL PI together with 2.5 mM DTDP, followed by time-lapse live imaging for 2 hours. **(B)** Time-lapse images show baseline-dead neurons become double-positive after PI addition (T = 0 min; yellow). Vulnerable neurons become PI-positive over time after DTDP exposure (T = 30 and 120 min; red), while resilient neurons remain non-fluorescent. Bottom panels show high-magnification views of the boxed regions in the upper panels. Scale bar, 50 μm. **(C)** Quantification of SG and PI fluorescence intensity over time for dead neurons (left panels) and live neurons (right panels) at T = 0 min. Average Mean ± Standard deviation (SD). N = 20 neurons.

We conducted quantification in which we created regions of interest (ROIs) on SG (+)-dead or SG (-)-living neurons at T = 0 min (i.e., right after DTDP + PI treatment) and measured SG and PI fluorescence intensity over time **(Fig. 2C)**. As expected, we found that baseline-dead neurons exhibit significantly higher SG and PI fluorescence compared to SG-negative live neurons upon PI + DTDP incubation. We also found that baseline SG-negative live neurons display increased PI fluorescence over time in response to DTDP. Of note, we observed that the SG signal unexpectedly decays over time in baseline-dead neurons, most likely due to dissociation of SG from DNA, which appears to contribute to a slightly increased SG signal in vulnerable neurons. Taken together, these data suggest that the sequential dual-dye incubation approach successfully labels baseline-dead, newly dying vulnerable neurons, and neurons resilient to oxidative stress in different colors within the same culture.

### DTDP induces dose-dependent acceleration of PI positivity and reveals cell-by-cell heterogeneity in neuronal vulnerability

To determine whether our cell-labeling approach can measure neuronal vulnerability in a gradual rather than an “on-off” manner, neurons were exposed to different concentrations of DTDP. Survival curve analysis of time-lapse imaging demonstrated a dose-dependent decrease in the probability of neurons remaining PI negative **(Fig. 3A)**. Moreover, the scatter plots of the same data sets revealed substantial cell-to-cell variability in the times becoming PI-positive, particularly at intermediate DTDP doses (e.g., 0.5-1 mM). These data support heterogeneous vulnerability and resilience states rather than a uniform response of individual neurons to mild oxidative stress and degeneration **(Fig. 3B)** and highlight the great sensitivity of our assay.

**Figure 3.**
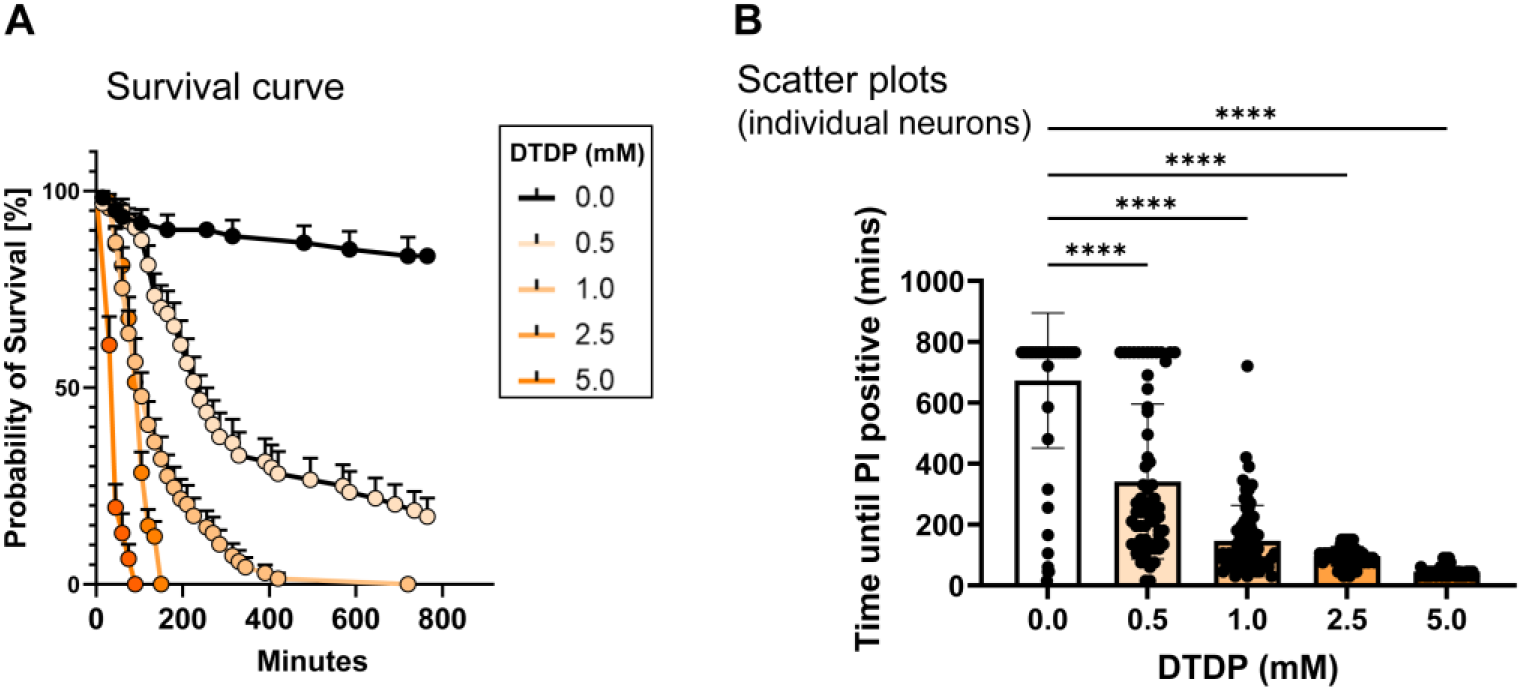
DTDP induces dose-dependent acceleration of PI positivity: **(A)** Survival curves showing the probability of survival (i.e., remaining PI-negative) over time following 5 μg/mL PI + DTDP treatment with different DTDP concentrations (0, 0.5, 1.0, 2.5, and 5.0 mM). **(B)** Bar plots of time until becoming PI-positive for individual neurons at each DTDP dose, with mean ± SEM. N = 46–74 neurons; Kruskal-Wallis test, ****p < 0.0001.

### Flow cytometry recapitulates subpopulations with distinct susceptibilities and defines a gating strategy

To further conduct mechanistic studies, we next aimed to physically separate and isolate subsets of neurons with different levels of vulnerability to DTDP-induced oxidative stress. As such, we adapted the sequential labeling workflow to flow cytometry and cell sorting (FACS). As shown in the experimental timeline **(Fig. 4A)**, neurons were incubated with 25 nM SG for 15 minutes, washed out the dye, and treated with 1 μg/mL PI in the presence of 0.5 mM DTDP or vehicle control for 30 minutes prior to flow cytometry analysis. To obtain a control containing only dead cells, we pre-incubated neurons with 1% Triton X for 10 minutes, then sequentially incubated with SG and PI. Surprisingly, fluorescence signals were not significantly different between DTDP and vehicle-treated neurons **(Fig. 4B)**. As one of several attempts to exclude noises and detect “pure” fluorescence signals from the main cell populations, we divided the entire signals into four different gates based on forward and side scatters **(Fig. 4C)**, followed by examining SG and PI fluorescence in the individual gates **(Fig. 4D)**. Strikingly, we found that one of the gating approaches (#4) reproducibly reveals three identical populations in vehicle-treated neurons; a 1^st^ population with SG and PI negative, a 2^nd^ that is with SG negative but PI positive (notably, they are slightly SG positive and this could be due to the SG dissociated from the DNA of baseline-dead neuron as described above), and a 3^rd^ that is with SG and PI positive. Importantly, Triton X pre-incubation eliminated the 1^st^ and 2^nd^ populations and revealed only the 3^rd^ population. Furthermore, DTDP treatment decreased the 1^st^ population while increasing the 2^nd^ population **(Fig. 4D)**.

**Figure 4.**
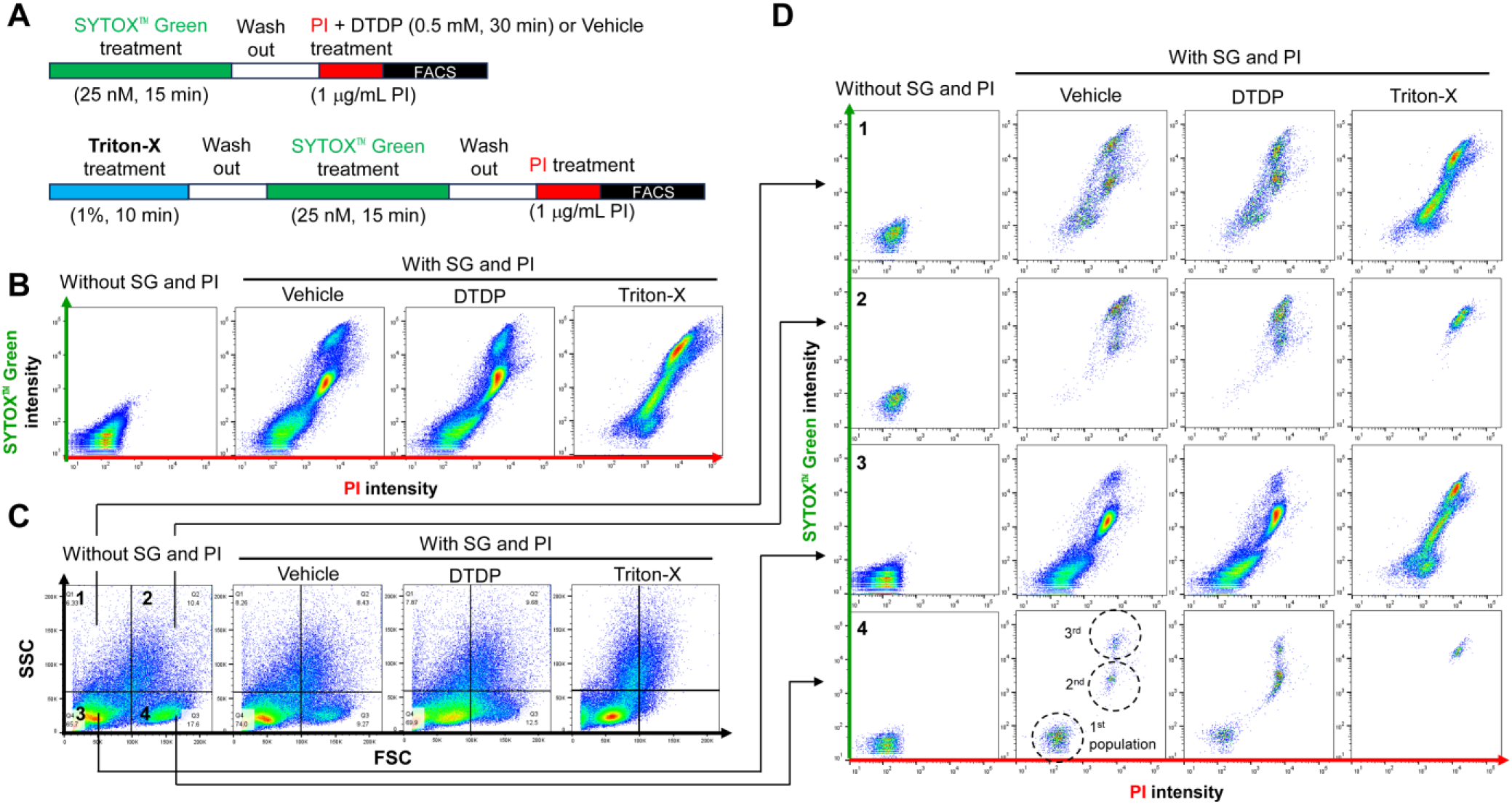
Flow cytometry gating strategy and controls for classifying vulnerable and resilient neuronal subpopulations to DTDP-induced oxidative stress and death: **(A)** Timeline for FACS experiments: neurons were treated with 25 nM SG for 15 min, washed, and then treated with 1 μg/mL PI plus DTDP (0.5 mM, 30 min) or vehicle prior to FACS. A positive-control using 1% Triton-X for 10 min incubation was included to define a dead-cell gate. **(B)** Representative SG versus PI bivariate plots illustrating how SG/PI quadrants shift under vehicle, DTDP, and Triton-X conditions without FSC/SSC gating. **(C)** Representative FSC/SSC plots showing the four different gates used to exclude debris and to select the neuronal population. **(D)** Representative SG versus PI bivariate plots under vehicle, DTDP, and Triton-X conditions when they are selectively analyzed across the indicated FSC/SSC gates (as linked by arrows). N = 5 independent experiments.

To fully validate our initial flow cytometry findings, we examined the dose-dependency of DTDP. After incubation of neurons with SG, and then washing out the medium, neurons were incubated with different concentrations of DTDP for 30 minutes in the presence of PI, and subjected to flow cytometry. In five independent experiments, we consistently found that the 1^st^ neuronal population (SG-/PI-) decreased, while the 2^nd^ population (SG-/PI+) increased in a dose-dependent manner **(Fig. 5A** and **5B)**. On the other hand, the number of cells in the 3^rd^ population (SG+/PI+) was not affected by different doses of DTDP **(Fig. 5A** and **5B)**. These results strongly suggest that 1^st^ and 2^nd^ populations correspond to resilient and vulnerable neurons, respectively. After sorting vulnerable and resilient neurons through 11 independent experiments (for the following lipidomic analysis), we validated cell viability using the Acridine Orange/Propidium Iodide Stain Kit. We found that resilient neurons exhibit significantly greater viability than vulnerable neurons **(Fig. 5C)**, further evidencing the successful separation and isolation of these two neuronal populations.

**Figure 5.**
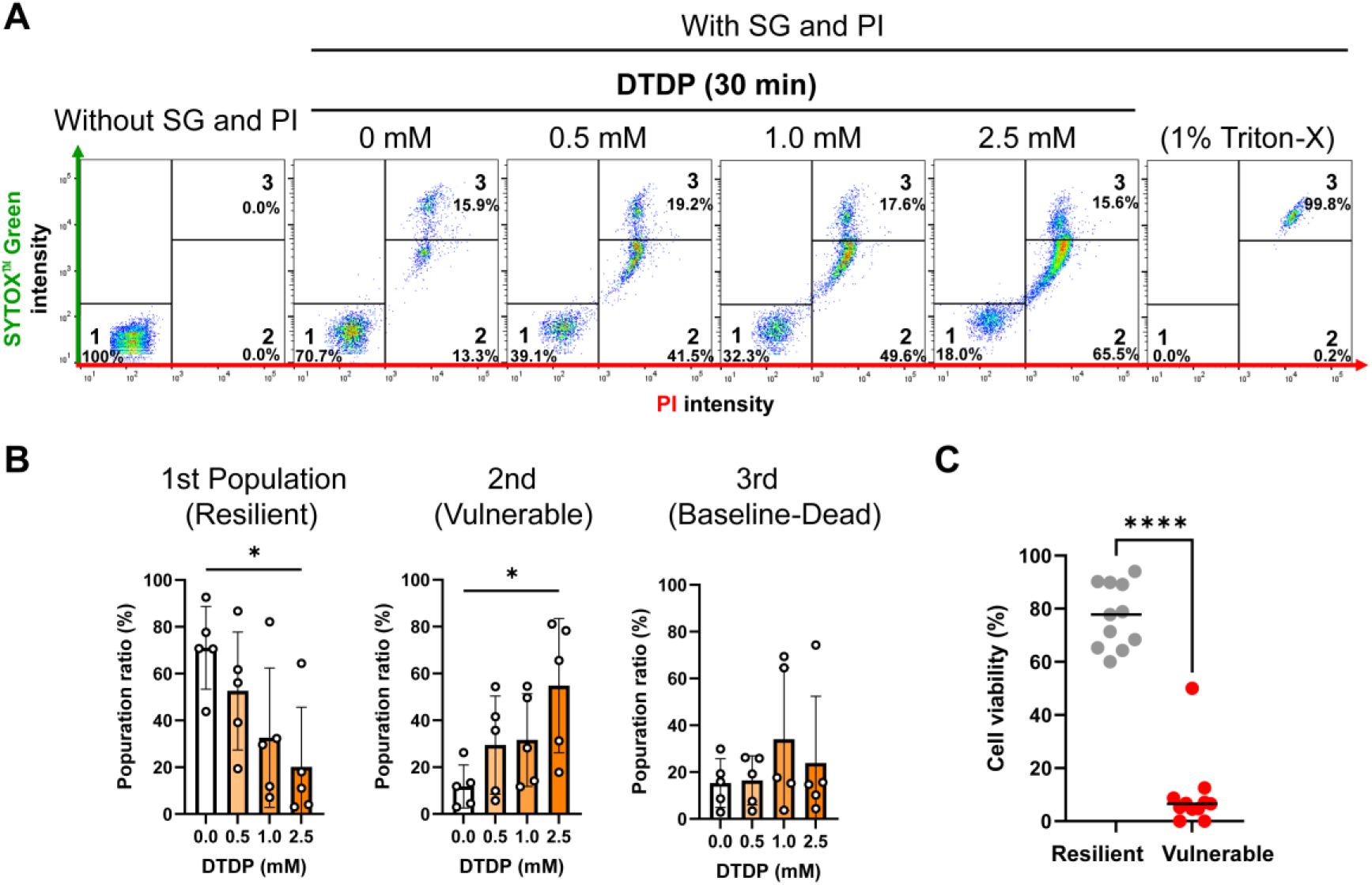
DTDP dose-dependent changes in FACS-defined neuronal populations: **(A)** Representative SG versus PI plots after 30 min DTDP treatment at the indicated doses (0, 0.5, 1.0, and 2.5 mM), with negative (without SG and PI) and 1% Triton-X positive controls. Quadrants define 1. SG−PI−: resilient, 2. SG−PI+: vulnerable, and 3. SG+/PI+: base-line dead neurons. **(B)** Quantifying the proportion of resilient, vulnerable, and baseline dead neurons across DTDP doses. N = 5 independent experiments; One-way ANOVA; *p < 0.05. **(C)** Cell viability assessment of sorted vulnerable and resilient neurons using the Acridine Orange/Propidium Iodide Stain Kit. N = 11 independent experiments; Paired Wilcoxon test, ****p<0.001.

### Membrane lipidomics identifies lipid-class features associated with neuronal vulnerability versus resilience to oxidative stress

Membrane permeability is directly assessed by our time-lapse imaging with PI. Moreover, lipid composition is one of the major determinants of membrane permeability by regulating the biophysical properties of the bilayer (Harayama and Riezman, 2018). Hence, we characterized the membrane lipid profiles of vulnerable and resilient neurons using lipidomic analysis as a “proof-of-concept” experiment to demonstrate how our new cell-sorting method can be adapted for mechanistic studies.

After sorting out the two populations of neurons treated with 1 mM DTDP for 30 minutes by FACS, the membrane fraction was extracted and subjected to LC–MS/MS-based lipidomic analysis (N = 11 independent experiments). In parallel, the number of neurons was measured using the Acridine Orange/Propidium Iodide Stain Kit. Immunoblotting validated successful fractionation and membrane enrichment, as indicated by Na^+^/K^+^-ATPase detection in the membrane fraction while β-tubulin enrichment in the cytoplasm **(Fig. 6, Supplemental Fig. 1)**. In the lipidomic analysis, the area size of the signal peak was measured for individual lipid species, and the values were normalized by the number of cells. Then, these values were converted to relative composition (%) by dividing the normalized signal of each lipid by the sum of all lipids to generate a percentage.

**Figure 6.**
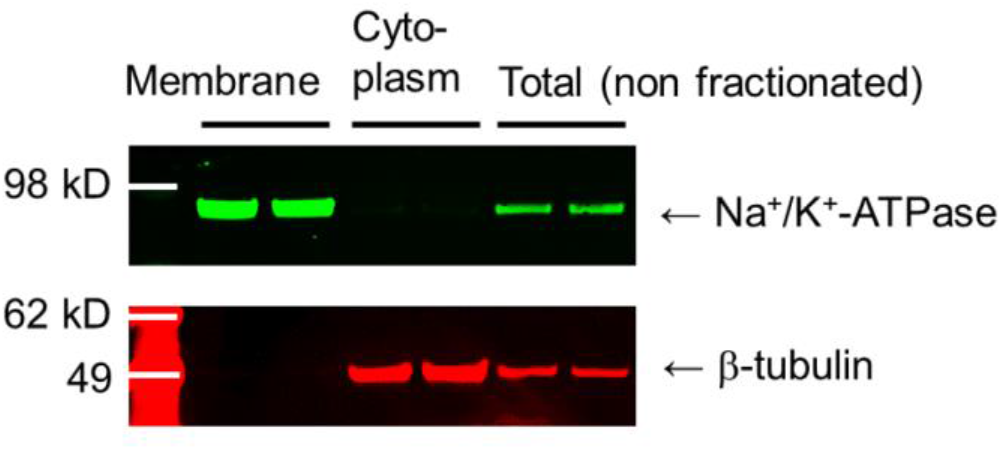
Extraction of membrane fraction: Western blotting validates successful fractionation using Na^+^/K^+^-ATPase (membrane marker) and β-tubulin (cytoplasmic marker) in membrane, cytoplasmic, and total (non-fractionated) samples.

First, we compared the membrane lipid class composition of vulnerable and resilient neurons to DTDP-induced oxidative stress, the heatmap of which is shown in **Fig. 7A**. Lipid classes were categorized into neutral lipids **(Fig. 7B),** sphingolipids **(Fig. 7C),** and phospholipids **(Fig. 7D),** and the relative abundance was visualized **(Fig. 7B-D, left panels)**. To further quantitatively assess differences in lipid classes between vulnerable and resilient neurons, the % difference, calculated as the relative ratio of the value in resilient over that in vulnerable neurons, was subjected to statistical analysis on a lipid class-by-class basis **(Fig. 7B-D, right panels)**. While majority of lipids are similarly expressed in the membranes of vulnerable and resilient neurons, some of the lipid classes significantly differed between the two neuronal populations, specifically ceramides (Cer) increased and monohexosylceramide (Hex1Cer) reduced in resilient compared to vulnerable neurons **(Fig. 7C)**. Individual lipid analysis identified Cer(d40:1)+H significantly increased in resilient neurons **(Fig. 7E)**. Of note, while lipid class analysis did not detect a statistical difference **(Fig. 7B** and **7D)**, there were several diacylglycerols (DG), triglycerides (TG), and phosphatidylcholines (PC) species that significantly differed between vulnerable and resilient neurons **(Fig. 7E)**. Lastly, we evaluated global saturation and chain length statuses between vulnerable and resilient neurons. To do so, lipids were categorized based on the total number of double bonds into saturated fatty acids (SFA; 0 double bonds), monounsaturated fatty acids (MUFA; 1 double bond), and polyunsaturated fatty acids (PUFA; 2 or more double bonds) **(Fig. 7F)**. Moreover, lipids were grouped by total acyl chain length into short-chain fatty acids (SCFA; <6 carbons), medium (MCFA; 6-12 carbons), long (LCFA; 13-21 carbons), and very long categories (VLCFA;>21 carbons) **(Fig. 7G)**. We found that global saturation and chain length statuses are not different between vulnerable and resilient neurons **(Fig. 7F** and **7G)**. Collectively, these results suggest that vulnerable and resilient neurons exhibit overall similar membrane lipid signatures; however, they differ significantly in some lipid components.

**Figure 7.**
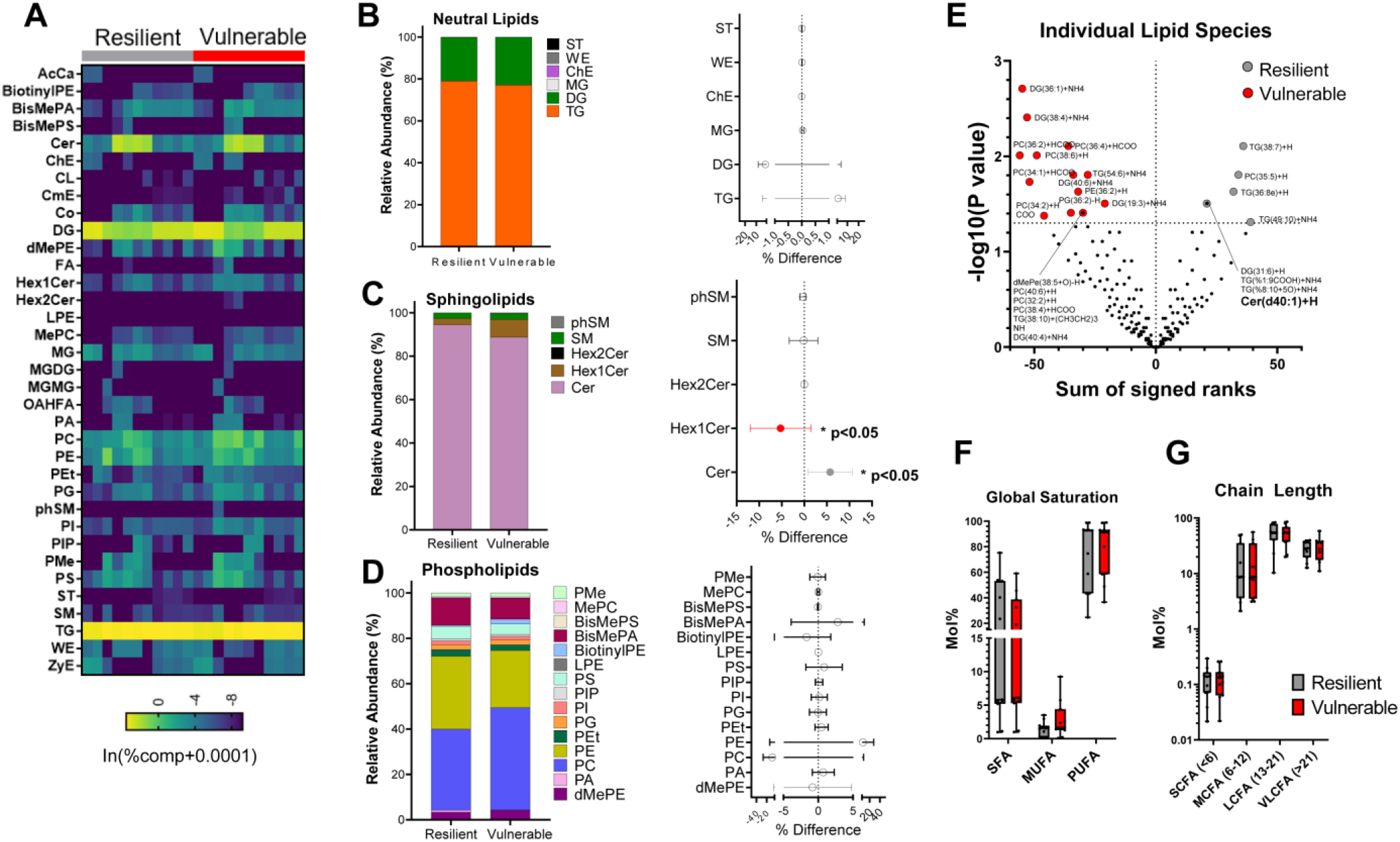
Membrane lipidomic analysis of vulnerable vs. resilient neurons to DTDP-induced oxidative stress and degeneration: Membrane fractions derived from FACS-isolated vulnerable and resilient neurons were subjected to LC–MS/MS lipidomic analysis (1 mM DTDP for 30 minutes) (N = 11 independent experiments). **(A)** Heatmap of relative lipid class abundance. **(B-D)** Left panels: lipid classes are categorized into neutral lipids, sphingolipids, and phospholipids, and the average relative abundance (%) in eleven experiments is displayed. Right panels: % Difference between vulnerable and resilient neurons. A positive % indicates lipid class increases in resilient neurons; on the other hand, a negative % means an increase in vulnerable neurons. Pairwise Wilcoxon tests, *p<0.05. **(E)** Volcano plots showing lipid species that were significantly increased in vulnerable (red dots) and resilient neurons (grey). Lastly, lipids are grouped by saturation status **(F)** and fatty acid length **(G)**, with no significant difference between vulnerable and resilient neurons.

### Time-lapse imaging establishes a cause-and-effect relationship between alterations in membrane lipids and neuronal vulnerability to DTDP-induced oxidative stress

Ceramides (Cer) and monohexosylceramide (Hex1Cer) increased in resilient and vulnerable neurons to DTDP-induced oxidative stress, respectively (**Fig. 7**); however, whether this alteration is a cause or a consequence remains elusive. To address this, we incubated neurons with 500 µM Conduritol B epoxide (CBE) for 5 days, a potent inhibitor of glucocerebrosidase (Gcase, a.k.a. GBA1), to prevent Hex1Cer metabolism into glucose and Cer (Grabowski et al., 1986) (**Fig. 8A**), followed by treatment with 0.5 mM DTDP. Using LC–MS/MS lipidomics, we confirmed that CBE increases the Hex1Cer over Cer ratio approximately 2.25-fold compared to vehicle control. Strikingly, we found that neurons treated with CBE degenerated significantly earlier than those with vehicle control in response to DTDP treatment (**Fig. 8B**). To ensure reproducibility, we conducted the same experiments twice more and found that one experiment exhibits similarly statistically different while another showing a consistent trend that neurons pre-incubated with CBE are more vulnerable to DTDP treatment than those with vehicle control (**Fig. 8B**). Moreover, we conducted survival curve analysis and found in all three independent experiments that the probability of survival is significantly decreased in neurons pre-incubated with CBE compared to vehicle control (**Supplemental Fig. 2A**). Next, we pre-incubated neurons with 10 µM Miglustat, an FDA-approved glucosylceramide synthase (GCS) inhibitor to block the conversion of Cer into Hex1Cer (Cox et al., 2000; Patterson et al., 2007) (**Fig. 8A**), or vehicle control for 5 days. Then, the neurons were treated with 0.5 mM DTDP to induce oxidative stress and neuronal loss. We found that neurons pre-incubated with Miglustat survive significantly longer than those treated with vehicle in the presence of DTDP (**Fig. 8C**).

**Figure 8.**
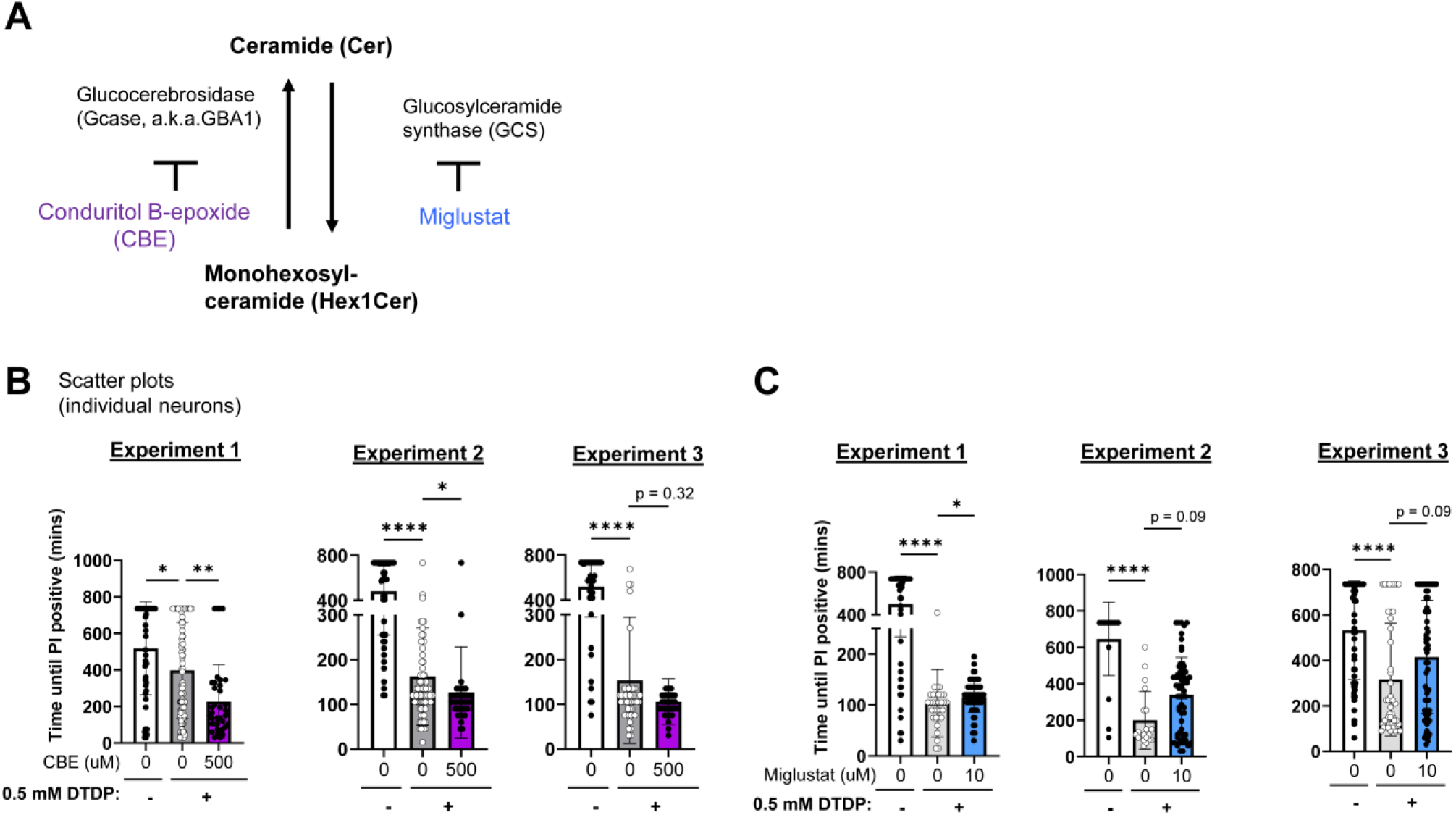
Cause-and-effect relationships between altered Cer-Hex1Cer axis and vulnerability to DTDP-induced oxidative stress and neuronal death: **(A)** Schematic of Cer and Hex1Cel conversion, responsible enzymes, and small compounds used in this study to modulate their activities. Neurons were treated with 500 μM CBE **(B)**, 10 μM Miglustat **(C)**, or vehicle control for 5-7 days, followed by time-lapse imaging to measure the time until becoming PI (+) on a cell-by-cell basis in response to 0.5 mM DTDP treatment. N = 3 independent experiments; One-way ANOVA or Kruskal-Wallis test; *p<0.05, **p<0.01, ****p<0.0001.

Although two other experiments did not reach a statistical difference, probably as there seems to be two drug responsive and resistant neuronal populations (plus, LC–MS/MS lipidomics verified that the Hex1Cer over Cer ratio decreases 1.33-fold only), we found a similar overall trend showing that neurons treated with Miglustat are more resilient to DTDP treatment (**Fig. 8C**). In survival curve analysis, we found in two out of three experiments that the probability of survival significantly increases in neurons pre-incubated with Miglustat compared to vehicle control (**Supplemental Fig. 2B**). Together, these results suggest that the Cer-Hex1Cer axis is causally associated with neuronal vulnerability to DTDP-induced oxidative stress and degeneration. However, it would be worth noting the absence of statistical difference in some experiments, as it suggests that 1) further careful investigation would be needed to modulate the Cer-Hex1Cer axis and 2) other factors, e.g., genes and proteins, could also be responsible for determining vulnerability and resilience to DTDP-induced degeneration.

### The link between the Cer-Hex1Cer axis and neuronal vulnerability is specific to DTDP-induced oxidative stress and degeneration

Under oxidative stress, reactive oxygen species (ROS) oxidize polyunsaturated fatty acids (PUFAs), leading to the formation of reactive aldehydes, including 4-Hydroxy-2-nonenal (HNE). HNE covalently binds not only to proteins, altering or inactivating their functions, but also to lipids such as phosphatidylethanolamine (PE) within the cell membrane (Guichardant et al., 1998). This alters lipid packing, reduces membrane bilayer thickness, and impairs membrane integrity (Hwang et al., 2008; Jovanović et al., 2019; Alonso et al., 2022). Importantly, HNE accumulates in the brains of various neurodegenerative diseases, such as Alzheimer’s disease (Markesbery and Lovell, 1998), Huntington disease (Lee et al., 2011), and ALS (Pedersen et al., 1998). Therefore, we used HNE to ask whether the involvement of Cer-Hex1Cer is DTDP-specific or is recapitulated by a mechanistically distinct stressor related to oxidative stress. As expected, we found that HNE shortens the time until becoming PI positive in a dose-dependent manner (**Fig. 9A**). Moreover, some neurons die while others survive in the presence of HNE (**Fig. 9A**), highlighting cell-by-cell heterogeneity in vulnerability to HNE treatment. We then isolated vulnerable and resilient neurons to HNE-induced degeneration by FACS, extracted the membrane fraction from sorted neurons, and subjected them to lipidomic analysis. We did not find significant differences in Ceramides (Cer) and monohexosylceramide (Hex1Cer) levels between resilient and vulnerable neurons to HNE treatment (**Fig. 9B**). To verify this using a different approach, we pre-incubated neurons with 500 µM CBE (**Fig. 9C**) or 10 µM Miglustat (**Fig. 9D**), then treated them with 200 µM HNE in the presence of PI for time-lapse imaging. In contrast to DTDP-treated neurons, CBE did not remarkably affect vulnerability to HNE-induced degeneration (**Fig. 9C and Supplemental Fig. 3A**). Moreover, neurons treated with Miglustat even tend to degenerate earlier compared to those with vehicle controls under HNE treatment (**Fig. 9D and Supplemental Fig. 3B**). Collectively, these results suggest that the Cer-Hex1Cer axis is selectively associated with neuronal vulnerability to DTDP-induced oxidative stress and degeneration.

**Figure 9.**
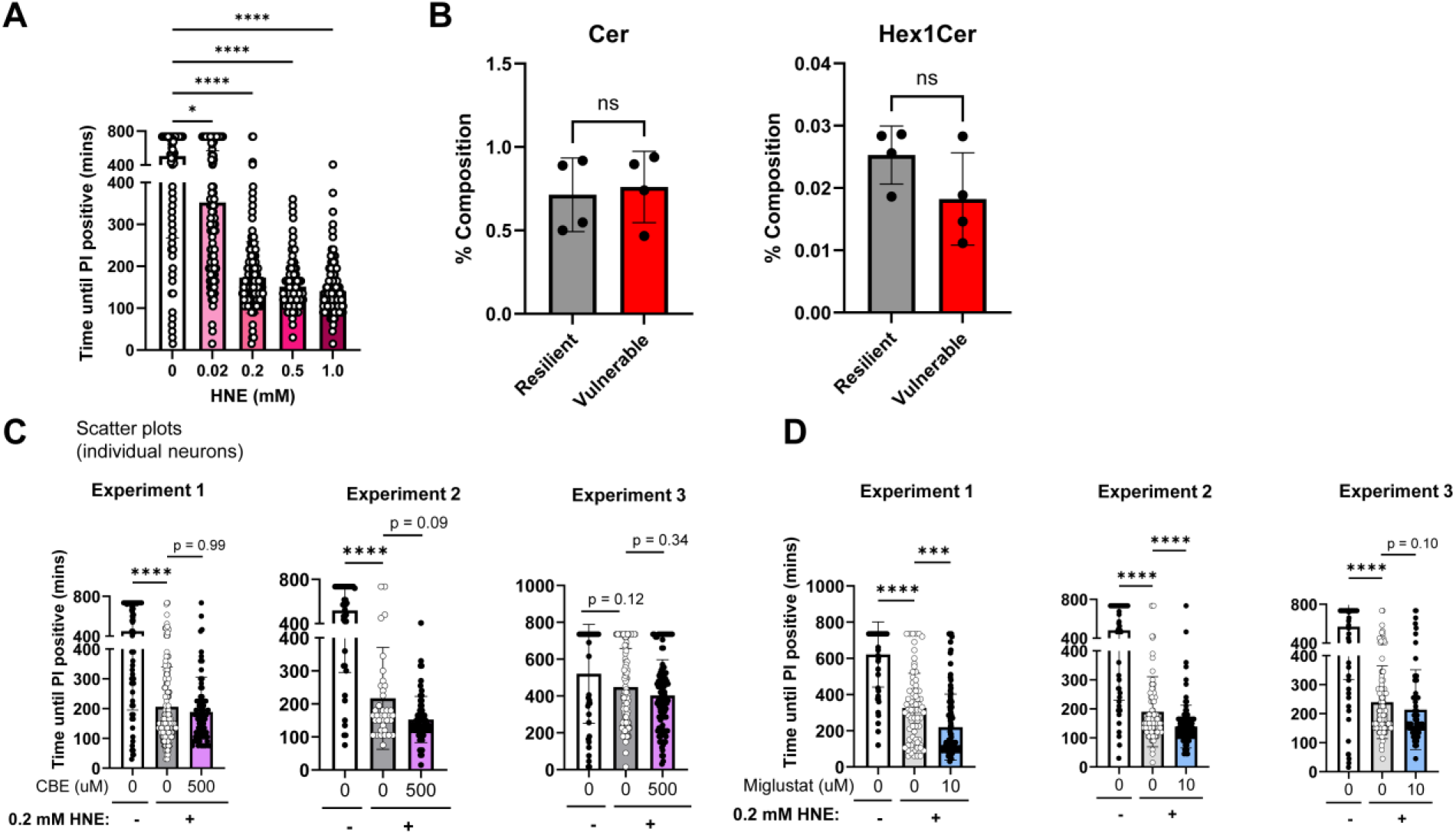
The Cer-Hex1Cer axis is not associated with neuronal vulnerability to HNE-induced degeneration: **(A)** Bar plots of time until becoming PI-positive for individual neurons at each HNE dose, with mean ± SEM. Kruskal-Wallis test, *p < 0.05, ****p < 0.0001. **(B)** LC-MS/MS lipidomics determined the relative abundance (%) of Cer and Hex1Cer in vulnerable and resilient neurons to 0.5 mM HNE treatment for 30 min. Time-lapse imaging of neurons pre-incubated with 500 μM CBE **(C)** or 10 μM Miglustat **(D)** to measure the time until becoming PI (+) on a cell-by-cell basis in response to 0.2 mM HNE treatment. N = 3 independent experiments; One-way ANOVA or Kruskal-Wallis test; n.s. no significance.

## Discussion

Some neurons die while others survive in response to toxic insults in neurodegenerative diseases. However, the mechanisms underlying the cell-by-cell heterogeneity in neuronal vulnerability to degeneration remain elusive. Using an oxidative stress paradigm as an example, this study reports the development of a novel methodology to identify and physically separate vulnerable from resilient neurons in culture. We employ a sequential cell-labeling workflow that uses two cell-impermeable fluorescence dyes (SYTOX™ Green [SG] and Propidium Iodide [PI]) (**Figs. 1** and **2**), which has been validated by time-lapse confocal imaging (**Figs. 2** and **3**), revealing vast cell-by-cell heterogeneity in neuronal vulnerability to DTDP-induced oxidative stress. To explore the factors associated with this heterogeneity, we subsequently used FACS to physically separate vulnerable from resilient neurons (**Figs. 4** and **5**) and subject them to mechanistic studies. As a proof-of-concept analysis, we have extracted the membrane fraction of isolated neurons (**Fig. 6**), subjected them to LC–MS/MS-based lipidomic analysis, and identified several lipid classes that are significantly different between vulnerable and resilient neurons (**Fig. 7**).

Selective neuronal vulnerability refers to the distinct susceptibilities of neuronal populations to cell injury and death in the context of neurodegenerative diseases. This phenomenon has been well documented across various brain regions and circuits, manifesting in stereotypical spatiotemporal patterns of cell death that are characteristic of distinct neurodegenerative diseases. For instance, AD involves the preferential loss of cholinergic neurons in the nucleus basalis of Meynert, noradrenergic neurons in the locus coeruleus, and neurons in the entorhinal cortex layers II, III (superficial), and IV (Kampmann 2024). Mechanistically, selective neuronal vulnerability is thought to be driven by a multitude of intrinsic neuronal and extrinsic factors: distribution of calcium-buffering proteins, metal transporters, antioxidant enzymes, and neurotransmitter receptors, differences in protein homeostatic factors (e.g., molecular chaperones and autophagy), differences in energy demand, varying susceptibilities to alterations in survival factor levels, and heterogeneity in glial interactions (Double et al., 2010; Kampmann 2024).

Recent advances in single-cell and spatial transcriptomics have enabled the systematic uncovering of selectively vulnerable neuronal populations from post-mortem brain tissue. For instance, Leng et al., identified and characterized RORB+ vulnerable neuronal subpopulations in the entorhinal cortex, finding upregulation of genes coding for potassium channels and downregulation of genes coding for pre- and post-synaptic proteins (Leng et al., 2021). Similarly, Lin et al., found a subset of phospho-tau (p-tau) vulnerable neurons expressing genes enriched for apoptotic pathways, cellular component disassembly, and regulation of vesicle-mediated transport (Lin et al., 2025). Moreover, Ehrenberg et al., showed that neurons of the locus coeruleus (LC), a brain region that shows early AD-related p-tau inclusions and neuron loss, upregulate cholesterol homeostatic genes and downregulate antioxidant metallothionein genes relative to neurons of the substantia nigra, which is relatively resistant to AD-related degeneration despite having a similar cytoarchitecture (Ehrenberg et al., 2025). Beyond scRNA-seq in human tissues, a study using the bacTRAP technology in mice found that neurons from vulnerable brain regions (i.e., CA1, EC layer II) highly express genes involved in calcium signaling, serotonin signaling, and response to growth factors, while neurons from resistant brain regions highly express genes involved in the regulation of synaptic plasticity, synaptic transmission, and projection development (Roussarie et al., 2020).

However, the research field has faced several drawbacks that limit our ability to fully elucidate the fundamental mechanistic drivers of selective neuronal vulnerability. Firstly, while post-mortem brain tissues can provide robust, physiologically relevant information, they do not allow us to distinguish primary drivers of selective vulnerability from downstream epiphenomena or consequences of neurodegeneration due to their cross-sectional natures. In light of these challenges, our group has developed an *in cellulo* neuronal system that enables the identification and separation of vulnerable and resilient neurons and their subsequent use for mechanistic studies. By introducing a defined toxic insult and longitudinal monitoring of cell death at the single cell level, our system captures the dynamic trajectory of neurodegeneration and enables precise definition of the upstream cellular and molecular properties associated with neuronal vulnerability. This aspect is further augmented by our model’s ability to discriminate already-dead neurons (that may have died due to a multitude of reasons in culture) from vulnerable neurons that die specifically in response to the toxic insult; hence, while post-mortem tissue offers a static snapshot of late-stage/end-stage neurodegeneration, our model offers a controlled framework to look at cellular responses immediately following injury.

There are several key implicit advantages of our model that remain to be explored and leveraged in future studies. First, our method allows us to study various functional properties potentially underlying selective vulnerability beyond membrane lipids, including genes and proteins. One of the other benefits is the prospect of characterizing drivers of selective vulnerability to distinct exogenous (e.g., oxidative stress, excitotoxicity, misfolded proteins, hypoxic injury) or endogenous (genetic background) stressors; this is crucial since different pathways most likely confer susceptibility to different toxic stimuli and intrinsic properties (Saxena et al., 2011). More broadly, our method can be tailored to a variety of cell types, including astrocytes and microglia, which have been shown to play a key role in selective neuronal vulnerability (Wang et al., 2010); this paves the way for elucidating a more comprehensive understanding of selective vulnerability that encompasses the complexities of neuron-glial functional interactions.

As a proof-of-concept analysis demonstrating that subsequent mechanistic analyses can be performed following cell isolation, we have characterized the membrane lipid signatures of vulnerable and resilient neurons to oxidative stress. This is because our time-lapse imaging assay is based on the quantitative assessment of membrane integrity, which is tightly associated with membrane lipid compositions (Harayama and Riezman, 2018). Intriguingly, our study identified significant differences in the membrane levels of two lipid classes including ceramide (Cer) and monohexosylceramide (Hex1Cer) (**Fig. 7**).

Generally, literature shows a positive association of ceramide (Cer) with neurodegeneration. For instance, Cer levels are elevated in the blood, cerebrospinal fluid (CSF), brain tissue, and adjacent to plaques in individuals with AD (Han et al., 2002; Mielke et al., 2010; Panchal et al., 2014; Teitsdottir et al., 2021). Similarly, a study in mice has found that Apo-E4 expression drives the upregulation of several Cer species in the cortex (Miranda et al., 2022). Mechanistically, Cer has been shown to promote amyloidogenesis through the stabilization of β-secretase as well as promote neuronal apoptosis (Jazvinšćak Jembrek et al., 2015). Nevertheless, it is also reported that Cer levels in cell membranes are upregulated in response to various insults, including tumor necrosis factor-alpha, UV and ionizing radiation, endotoxin, and chemotherapeutic agents; Zhao et al. show that the addition of ceramides to lipid monolayers actually stiffens the membrane packing and reduce its permeability, thus potentially acting to reduce cellular damage from chemicals (e.g., hydroxyl free radicals) (Zhao et al., 2022). Thus, our finding that Cer levels are elevated in resilient neurons is consistent with this potentially protective role, whereby neurons that are able to upregulate Cer or have higher levels at baseline in their membranes are more likely to survive even in the presence of toxic insults. Moreover, Cer and acid sphingomyelinase are known to be actively involved in the repair process of damaged plasma membrane (Tam et al., 2010). This also raises the possibility that our resilient neurons may exhibit a greater capacity for plasma membrane repair than vulnerable neurons, as demonstrated by increased Cer levels.

In contrast, the functional link between monohexosylceramide (Hex1Cer) and neuronal survival has been well established. This is because homozygous mutations in the lysosomal enzyme glucocerebrosidase (GCase, a.k.a. GBA1), hydrolyzing Hex1Cer, cause an inherited lysosomal storage disease known as Gaucher disease (Tsuji et al., 1987), which involves severe neurodegeneration. In the same line, GBA1 deficient mice exhibit severe neurodegenerative phenotypes (Enquist et al., 2007). In addition, heterozygous mutations in GBA1 are also the major known genetic risk factor for Parkinson’s disease (Sidransky et al., 2009). Miglustat, a small molecule inhibitor of glucosylceramide synthase, has been used to treat lysosomal storage disorders in clinical settings (Cox et al., 2000; Patterson et al., 2007). Indeed, our data showed that neurons treated with Miglustat are likely more resistant to DTDP-induced neuronal death, whereas CBE, a potent inhibitor of GCase, shows the opposite effect (**Fig. 8**). Interestingly, this effect is likely specific to DTDP, perhaps linked to general lipid peroxidation, since HNE-induced membrane permeabilization is not meaningfully affected by CBE or Miglustat treatment (**Fig. 9**) compared to DTDP (**Fig. 8**). Together, these findings suggest that Hex1Cer could be one of the factors regulating neuronal resilience to oxidative stress, particularly in the context of lipid peroxidation-mediated membrane permeabilization.

Despite the promise of our methodology, several limitations persist. First, our current methodology does not allow us to isolate vulnerable neurons at the exact moment of their death, but rather shortly after; thus, certain protein and RNA species may be affected by increased membrane permeability, which could affect their quantification. Moreover, our model has been tested only on mouse primary neurons; whether human iPSC-derived neurons are compatible with the sequential staining method and subsequent FACS needs to be tested. Lastly, our method is based on 2D mono-culture neurons, which lack an *in vivo* cellular context. This could be crucial since glial cells, particularly astrocytes, play a critical role in neuronal lipid metabolism (Barber and Raben, 2019).

In summary, we report the development of a novel methodology to identify and isolate vulnerable and resilient neurons in culture. This utilizes a sequential dual-dye incubation approach to differentially label cells based on the integrity of their membranes after the introduction of oxidative stress, followed by FACS to physically separate and isolate the two neuron subsets with different susceptibilities. As a proof-of-concept analysis of a subsequent mechanistic investigation, we used this isolation method to characterize the membrane lipid signatures of vulnerable and resilient neurons, finding that ceramide (Cer) and monohexosylceramide (Hex1Cer) levels differ significantly between the two cell populations. Overall, this methodology lays the foundation for a controlled, mechanistic study of the cellular and molecular properties that underlie selective neuronal vulnerability, paving the way for the identification of therapeutic targets and diagnostic markers for neurodegenerative diseases.

## Acknowledgements

We thank Dr. John Asara (BIDMC Mass Spectrometry Core) for lipidomic analysis, and Drs. Alberto Serrano-Pozo, Steven Hou, and Bradley Hyman (MGH Neurology) for constructive feedback on the manuscript. This work was funded by the National Institutes of Health grants AG079838 (to M. M.).

## Author contributions

Conceptualization: TGI, MM; Data curation: MS, MY; Formal analysis: MS, MY; Funding acquisition: MM, Investigation: MS, MY; Methodology: DOK, REB; Project administration: MM, Resources: EW, NW; Software: MS, MY; Supervision: MM; Validation: MS, MY, SD; Visualization: MS, MY; Writing – original draft: MS, MY; Writing – review & editing: SD, REB, TGI, MM

## Declaration of interests

The authors declare no competing interests.

## Notes

### Competing Interest Statement

The authors have declared no competing interest.

## References

Alonso L, Menegatti R, Dorta ML, Alonso A. Plasma membrane rigidity effects of 4-hydroxy-2-nonenal in Leishmania, erythrocyte and macrophage. Toxicol In Vitro. 2022 Mar;79:105294. doi: 10.1016/j.tiv.2021.105294. Epub 2021 Dec 9. PMID: 34896601.

Andreone, B.J., Larhammar, M., and Lewcock, J.W. (2020). Cell Death and Neurodegeneration. Cold Spring Harb Perspect Biol 12, a036434. 10.1101/cshperspect.a036434.

Barber, C.N., and Raben, D.M. (2019). Lipid Metabolism Crosstalk in the Brain: Glia and Neurons. Front Cell Neurosci 13, 212. 10.3389/fncel.2019.00212.

Breitkopf, S.B., Ricoult, S.J.H., Yuan, M., Xu, Y., Peake, D.A., Manning, B.D., and Asara, J.M. (2017). A relative quantitative positive/negative ion switching method for untargeted lipidomics via high resolution LC-MS/MS from any biological source. Metabolomics 13, 30. 10.1007/s11306-016-1157-8.

Cox T, Lachmann R, Hollak C, Aerts J, van Weely S, Hrebícek M, Platt F, Butters T, Dwek R, Moyses C, Gow I, Elstein D, Zimran A. Novel oral treatment of Gaucher’s disease with N-butyldeoxynojirimycin (OGT 918) to decrease substrate biosynthesis. Lancet. 2000 Apr 29;355(9214):1481–5. doi: 10.1016/S0140-6736(00)02161-9. PMID: 10801168.

Double, K.L., Reyes, S., Werry, E.L., and Halliday, G.M. (2010). Selective cell death in neurodegeneration: Why are some neurons spared in vulnerable regions? Progress in Neurobiology 92, 316–329. 10.1016/j.pneurobio.2010.06.001.

Ehrenberg AJ, Sant C, Pereira FL, Li SH, Buxton J, Langlois S, Trinidad M, Oh I, Leite REP, Rodriguez RD, Paes VR, Pasqualucci CA, Seeley WW, Spina S, Suemoto CK, Temple S, Kaufer D, Grinberg LT. Pathways underlying selective neuronal vulnerability in Alzheimer’s disease: Contrasting the vulnerable locus coeruleus to the resilient substantia nigra. Alzheimers Dement. 2025 Mar;21(3):e70087. doi: 10.1002/alz.70087.

Enquist IB, Lo Bianco C, Ooka A, Nilsson E, Månsson JE, Ehinger M, Richter J, Brady RO, Kirik D, Karlsson S. Murine models of acute neuronopathic Gaucher disease. Proc Natl Acad Sci U S A. 2007 Oct 30;104(44):17483–8. doi: 10.1073/pnas.0708086104. Epub 2007 Oct 22. PMID: 17954912; PMCID: PMC2077282.

Fu, H., Hardy, J., and Duff, K.E. (2018). Selective vulnerability in neurodegenerative diseases. Nat Neurosci 21, 1350–1358. 10.1038/s41593-018-0221-2.

Grabowski GA, Osiecki-Newman K, Dinur T, Fabbro D, Legler G, Gatt S, Desnick RJ. Human acid beta-glucosidase. Use of conduritol B epoxide derivatives to investigate the catalytically active normal and Gaucher disease enzymes. J Biol Chem. 1986 Jun 25;261(18):8263–9. PMID: 3087971.

Guichardant M, Taibi-Tronche P, Fay LB, Lagarde M. Covalent modifications of aminophospholipids by 4-hydroxynonenal. Free Radic Biol Med. 1998 Dec;25(9):1049–56. doi: 10.1016/s0891-5849(98)00149-x. PMID: 9870558.

Han, X., M Holtzman, D., McKeel, D.W., Kelley, J., and Morris, J.C. (2002). Substantial sulfatide deficiency and ceramide elevation in very early Alzheimer’s disease: potential role in disease pathogenesis. J Neurochem 82, 809–818. 10.1046/j.1471-4159.2002.00997.x.

Harayama T, Riezman H. Understanding the diversity of membrane lipid composition. Nat Rev Mol Cell Biol. 2018 May;19(5):281–296. doi: 10.1038/nrm.2017.138. Epub 2018 Feb 7. Erratum in: Nat Rev Mol Cell Biol. 2019 Nov;20(11):715. doi: 10.1038/s41580-019-0171-x. PMID: 29410529.

Hwang JJ, Lee SJ, Kim TY, Cho JH, Koh JY. Zinc and 4-hydroxy-2-nonenal mediate lysosomal membrane permeabilization induced by H2O2 in cultured hippocampal neurons. J Neurosci. 2008 Mar 19;28(12):3114–22. doi: 10.1523/JNEUROSCI.0199-08.2008. PMID: 18354014; PMCID: PMC6670692.

Jazvinšćak Jembrek, M., Hof, P.R., and Šimić, G. (2015). Ceramides in Alzheimer’s Disease: Key Mediators of Neuronal Apoptosis Induced by Oxidative Stress and Aβ Accumulation. Oxid Med Cell Longev 2015, 346783. 10.1155/2015/346783.

Jovanović O, Škulj S, Pohl EE, Vazdar M. Covalent modification of phosphatidylethanolamine by 4-hydroxy-2-nonenal increases sodium permeability across phospholipid bilayer membranes. Free Radic Biol Med. 2019 Nov 1;143:433–440. doi: 10.1016/j.freeradbiomed.2019.08.027. Epub 2019 Aug 25. PMID: 31461663; PMCID: PMC7115857.

Kampmann, M. (2024). Molecular and cellular mechanisms of selective vulnerability in neurodegenerative diseases. Nat. Rev. Neurosci. 25, 351–371. 10.1038/s41583-024-00806-0.

Lee J, Kosaras B, Del Signore SJ, Cormier K, McKee A, Ratan RR, Kowall NW, Ryu H. Modulation of lipid peroxidation and mitochondrial function improves neuropathology in Huntington’s disease mice. Acta Neuropathol. 2011 Apr;121(4):487–98. doi: 10.1007/s00401-010-0788-5. Epub 2010 Dec 16. PMID: 21161248; PMCID: PMC3151672.

Leng K, Li E, Eser R, Piergies A, Sit R, Tan M, Neff N, Li SH, Rodriguez RD, Suemoto CK, Leite REP, Ehrenberg AJ, Pasqualucci CA, Seeley WW, Spina S, Heinsen H, Grinberg LT, Kampmann M. Molecular characterization of selectively vulnerable neurons in Alzheimer’s disease. Nat Neurosci. 2021 Feb;24(2):276–287. doi: 10.1038/s41593-020-00764-7.

Lin G, Chancellor SE, Kwon T, Woodbury ME, Doering A, Abdourahman A, Bennett RE, Liao F, Pastika T, Tamm J, Romanul N, Yanamandra K, Hu M, Zhao K, Frosch MP, Grinberg Y, Li H, Das S, Dellovade T, Karran EH, Talanian RV, Biber K, Serrano-Pozo A, Ried JS, Langlois X, Hyman BT. Cell-death pathways and tau-associated neuronal vulnerability in Alzheimer’s disease. Cell Rep. 2025 Jun 24;44(6):115758. doi: 10.1016/j.celrep.2025.115758.

Lundin, B., Wieckiewicz, N., Yokomizo, M., Sadek, M., Kwarteng, D.O., Dickson, J.R., Sobolewski, R.G.R., Derosla, V., Armagan, G., Perrin, F., et al. (2025). APP family is a regulator of endo-lysosomal membrane vulnerability. J Biol Chem 301, 110774. 10.1016/j.jbc.2025.110774.

Markesbery WR, Lovell MA. Four-hydroxynonenal, a product of lipid peroxidation, is increased in the brain in Alzheimer’s disease. Neurobiol Aging. 1998 Jan-Feb;19(1):33–6. doi: 10.1016/s0197-4580(98)00009-8. PMID: 9562500.

Mielke, M.M., Haughey, N.J., Bandaru, V.V.R., Schech, S., Carrick, R., Carlson, M.C., Mori, S., Miller, M.I., Ceritoglu, C., Brown, T., et al. (2010). Plasma ceramides are altered in mild cognitive impairment and predict cognitive decline and hippocampal volume loss. Alzheimers Dement 6, 378–385. 10.1016/j.jalz.2010.03.014.

Miranda, A.M., Ashok, A., Chan, R.B., Zhou, B., Xu, Y., McIntire, L.B., Area-Gomez, E., Di Paolo, G., Duff, K.E., Oliveira, T.G., et al. (2022). Effects of APOE4 allelic dosage on lipidomic signatures in the entorhinal cortex of aged mice. Transl Psychiatry 12, 129. 10.1038/s41398-022-01881-6.

Moujalled, D., Strasser, A., and Liddell, J.R. (2021). Molecular mechanisms of cell death in neurological diseases. Cell Death Differ 28, 2029–2044. 10.1038/s41418-021-00814-y.

Panchal, M., Gaudin, M., Lazar, A.N., Salvati, E., Rivals, I., Ayciriex, S., Dauphinot, L., Dargère, D., Auzeil, N., Masserini, M., et al. (2014). Ceramides and sphingomyelinases in senile plaques. Neurobiology of Disease 65, 193–201. 10.1016/j.nbd.2014.01.010.

Patterson MC, Vecchio D, Prady H, Abel L, Wraith JE. Miglustat for treatment of Niemann-Pick C disease: a randomised controlled study. Lancet Neurol. 2007 Sep;6(9):765–72. doi: 10.1016/S1474-4422(07)70194-1. PMID: 17689147.

Pedersen WA, Fu W, Keller JN, Markesbery WR, Appel S, Smith RG, Kasarskis E, Mattson MP. Protein modification by the lipid peroxidation product 4-hydroxynonenal in the spinal cords of amyotrophic lateral sclerosis patients. Ann Neurol. 1998 Nov;44(5):819–24. doi: 10.1002/ana.410440518. PMID: 9818940.

Roussarie JP, Yao V, Rodriguez-Rodriguez P, Oughtred R, Rust J, Plautz Z, Kasturia S, Albornoz C, Wang W, Schmidt EF, Dannenfelser R, Tadych A, Brichta L, Barnea-Cramer A, Heintz N, Hof PR, Heiman M, Dolinski K, Flajolet M, Troyanskaya OG, Greengard P. Selective Neuronal Vulnerability in Alzheimer’s Disease: A Network-Based Analysis. Neuron. 2020 Sep 9;107(5):821–835.e12. doi: 10.1016/j.neuron.2020.06.010.

Saura, C.A., Deprada, A., Capilla-López, M.D., and Parra-Damas, A. (2023). Revealing cell vulnerability in Alzheimer’s disease by single-cell transcriptomics. Seminars in Cell & Developmental Biology 139, 73–83. 10.1016/j.semcdb.2022.05.007.

Saxena, S., and Caroni, P. (2011). Selective Neuronal Vulnerability in Neurodegenerative Diseases: from Stressor Thresholds to Degeneration. Neuron 71, 35–48. 10.1016/j.neuron.2011.06.031.

Sidransky E, Nalls MA, Aasly JO, Aharon-Peretz J, Annesi G, Barbosa ER, Bar-Shira A, Berg D, Bras J, Brice A, Chen CM, Clark LN, Condroyer C, De Marco EV, Dürr A, Eblan MJ, Fahn S, Farrer MJ, Fung HC, Gan-Or Z, Gasser T, Gershoni-Baruch R, Giladi N, Griffith A, Gurevich T, Januario C, Kropp P, Lang AE, Lee-Chen GJ, Lesage S, Marder K, Mata IF, Mirelman A, Mitsui J, Mizuta I, Nicoletti G, Oliveira C, Ottman R, Orr-Urtreger A, Pereira LV, Quattrone A, Rogaeva E, Rolfs A, Rosenbaum H, Rozenberg R, Samii A, Samaddar T, Schulte C, Sharma M, Singleton A, Spitz M, Tan EK, Tayebi N, Toda T, Troiano AR, Tsuji S, Wittstock M, Wolfsberg TG, Wu YR, Zabetian CP, Zhao Y, Ziegler SG. Multicenter analysis of glucocerebrosidase mutations in Parkinson’s disease. N Engl J Med. 2009 Oct 22;361(17):1651–61. doi: 10.1056/NEJMoa0901281. PMID: 19846850; PMCID: PMC2856322.

Tam, C., Idone, V., Devlin, C., Fernandes, M.C., Flannery, A., He, X., Schuchman, E., Tabas, I., and Andrews, N.W. (2010). Exocytosis of acid sphingomyelinase by wounded cells promotes endocytosis and plasma membrane repair. J Cell Biol 189, 1027–1038. 10.1083/jcb.201003053.

Teitsdottir, U.D., Halldorsson, S., Rolfsson, O., Lund, S.H., Jonsdottir, M.K., Snaedal, J., and Petersen, P.H. (2021). Cerebrospinal Fluid C18 Ceramide Associates with Markers of Alzheimer’s Disease and Inflammation at the Pre- and Early Stages of Dementia. J Alzheimers Dis 81, 231–244. 10.3233/JAD-200964.

Tsuji S, Choudary PV, Martin BM, Stubblefield BK, Mayor JA, Barranger JA, Ginns EI. A mutation in the human glucocerebrosidase gene in neuronopathic Gaucher’s disease. N Engl J Med. 1987 Mar 5;316(10):570–5. doi: 10.1056/NEJM198703053161002. PMID: 2880291.

Wang, X., and Michaelis, E.K. (2010). Selective neuronal vulnerability to oxidative stress in the brain. Front. Aging Neurosci. 2. 10.3389/fnagi.2010.00012.

Yokomizo, M., Sadek, M., Williams, E., Houser, M.C.Q., Wieckiewicz, N., Torres, S., Berezovska, O., and Maesako, M. (2025). A unique subpopulation of wild-type neurons recapitulating familial Alzheimer’s disease phenotypes. Cell Death Dis 16, 604. 10.1038/s41419-025-07934-0.

Zhao, L., Liao, C., Chen, D., Zhang, D., Li, G., and Zhang, X. (2022). Stiffening Effect of Ceramide on Lipid Membranes Provides Non-Sacrificial Protection against Potent Chemical Damage. Langmuir 38, 3522– 3529. 10.1021/acs.langmuir.1c03427.

