## Supplemental Figures for "Separation, isolation, and membrane lipid profiling of selectively vulnerable and resilient neurons"

[#]These authors equally contributed

### Supplemental Figure 1

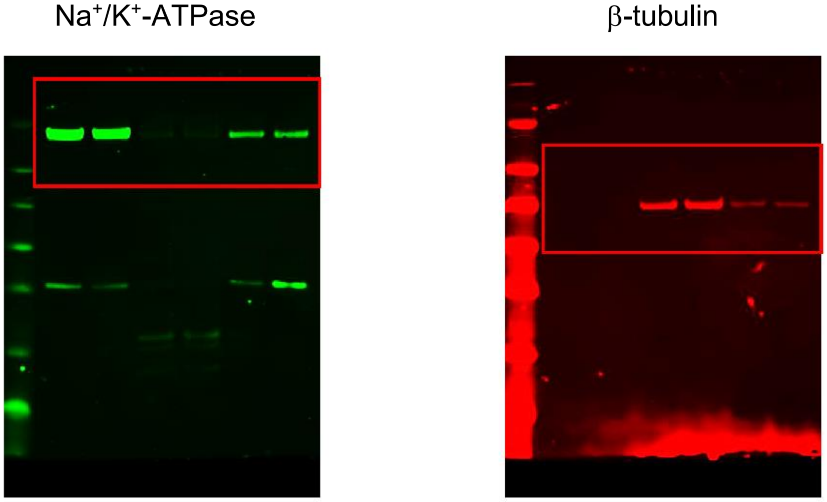


Raw Western Blot images of Figure 6.

### Supplemental Figure 2

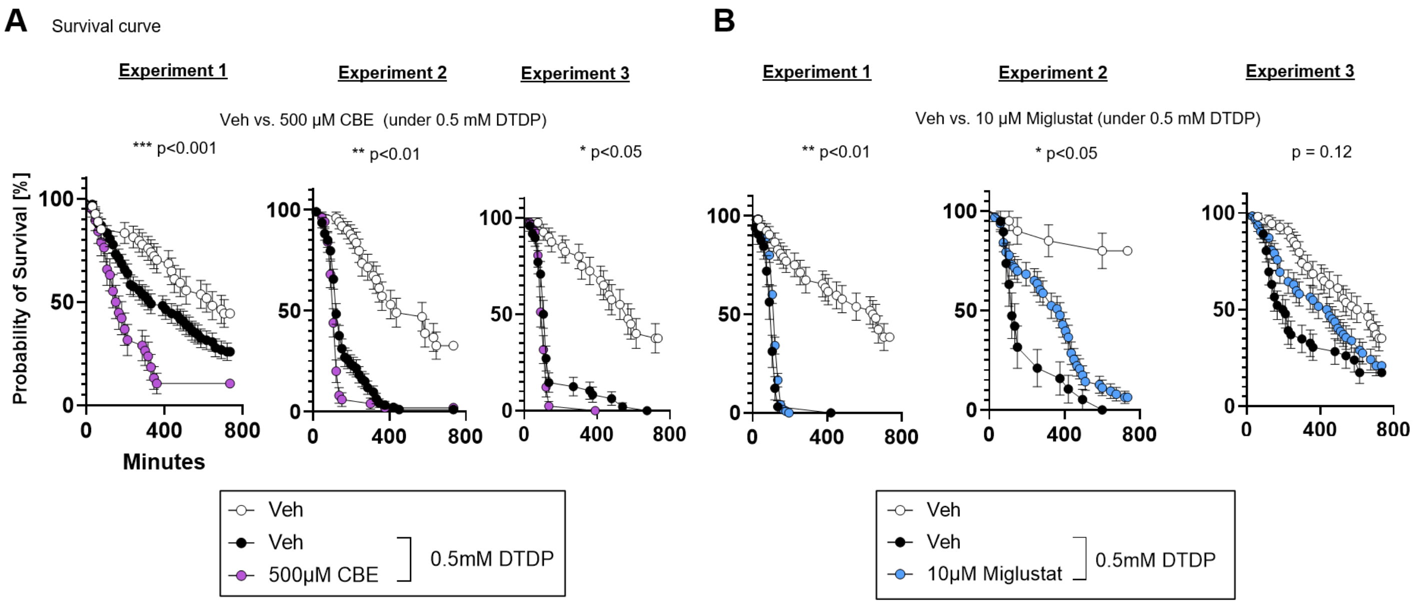


Survival curve analysis of Figure 8. Logrank (Mantel-Cox) test. *p<0.05, **p<0.01, ***p<0.001

### Supplemental Figure 3

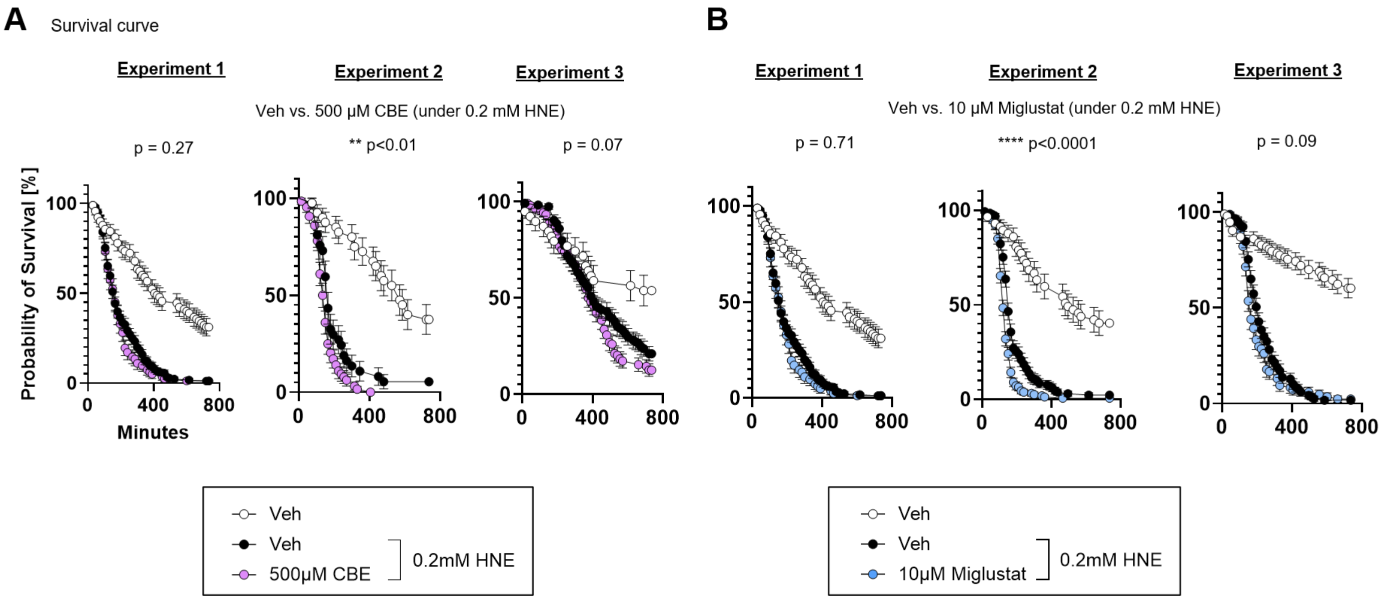


Survival curve analysis of Figure 9. Logrank (Mantel-Cox) test. **p<0.01, ****p<0.0001
